# Follicle-innervating Aδ-low threshold mechanoreceptors organize through a population-dependent mechanism

**DOI:** 10.1101/2022.08.09.503379

**Authors:** Matthew B. Pomaville, Kevin M. Wright

## Abstract

The mammalian somatosensory system is comprised of multiple neuronal populations that form specialized, highly organized sensory endings in the skin. The organization of somatosensory endings is essential to their functions, yet the mechanisms which regulate this organization remain unclear. Using a combination of genetic and molecular labeling approaches, we examined the development of mouse hair follicle-innervating low-threshold mechanoreceptors (LTMRs) and explored competition for innervation targets as a mechanism involved in the patterning of their receptive fields. We show that follicle innervating neurons are present in the skin at birth and that LTMR receptive fields gradually add follicle-innervating endings during the first two postnatal weeks. Using a constitutive *Bax* knockout to increase the number of neurons in adult animals, we show that two LTMR subtypes have differential responses to an increase in neuronal population size: Aδ-LTMR neurons shrink their receptive fields to accommodate the increased number of neurons innervating the skin, while C-LTMR neurons do not. Our findings suggest that competition for hair follicles to innervate plays a role in the patterning and organization of follicle-innervating LTMR neurons.

**Summary Statement:** Aδ follicle-innervating low-threshold mechanoreceptor neurons form tiled receptive fields through competition for hair follicles during the early postnatal period.

## Introduction

Mammalian somatosensation relies on the proper development, organization, and integration of multiple highly specialized sensory neuron subtypes that reside in the dorsal root ganglia (DRG) (Rice and Albrecht, 2008). In mice, there are at least 10 identified subtypes of skin-innervating sensory neurons, which can be broadly categorized as hairy skin-innervating or glabrous skin-innervating based on their terminal ending locations (Burgess et al., 1968; Cain et al., 2001; Halata, 1993; Iggo and Muir, 1969; Joong Woo Leem et al., 1993; Knibestöl, 1973; Lynn and Carpenter, 1982; Paré et al., 2002). Historically, these two groups of somatosensory neurons were subdivided based on their conduction velocity and responses to stimuli as measured by electrophysiological recordings (Brown and Iggo, 1967; Horch et al., 1977; Joong Woo Leem et al., 1993; Lewin and McMahon, 1991). In hairy skin, follicles are innervated by multiple populations of low-threshold mechanoreceptive neurons (follicle-innervating LTMRs).

Each LTMR neuron extends a peripherally projecting axon into the skin, where it elaborates a receptive field containing highly specialized endings around hair follicles. A series of landmark studies identified tools to genetically label somatosensory neuron subtypes, including the Aβ rapidly adapting (RA)-LTMR, Aδ-LTMR, and C-LTMR neurons, populations that were previously distinguished only by their electrophysiological characteristics (Li et al., 2011; Luo et al., 2009; Rutlin et al., 2014; Wu et al., 2012). These tools have enabled the examination of LTMRs on a population specific and single neuron level, allowing for more detailed questions about receptive field patterning and organization mechanisms to be asked.

In adult mice, the follicle-innervating LTMR neurons form discrete and stereotyped receptive fields, innervating a constrained number of hair follicles in the skin (Bai et al., 2015). LTMR subtypes show remarkable selectivity in the populations of hair follicles that they innervate. Aδ-LTMRs and C-LTMRs innervate both zigzag hairs and awl/auchene hairs, which comprise approximately 74% and 25% of the total hairs in the skin, respectively. Aβ RA-LTMRs form receptive fields innervating awl/auchene hairs, as well as guard hairs that comprise approximately 1% of the hairs in the skin (Kuehn et al., 2019; Li et al., 2011). Detailed electron microscopy studies also illustrated the ultrastructure of the longitudinal lanceolate ending (LLE), its ensheathing terminal Schwann cells, and their interaction with the hair follicle (Li and Ginty, 2014). These studies together laid the groundwork for molecularly driven exploration into the development and function of follicle-innervating neurons.

The strict organization of follicle innervating neurons into constrained receptive fields is essential for the proper localization of touch stimulus on the body. Recently, advancements in genetically driven multi-fluorophore reporter systems made it possible to examine the interactions between neurons of the same genetic subtype in adult mice. These studies showed that Aδ-LTMR and C-LTMR neurons form tiled receptive fields with minimal overlap between neighboring homotypic neurons. In contrast, Aβ RA-LTMR neurons form receptive fields that overlap with those of neighboring Aβ RA-LTMR neurons, resulting in LLEs from multiple Aβ RA-LTMR neurons at each guard hair (Kuehn et al., 2019). The mechanisms by which follicle-innervating LTMR neurons form exclusive receptive fields remains largely unexplored.

The homotypically tiled arrangement of Aδ-LTMRs and C-LTMRs in mice is reminiscent of the innervation patterns of dendritic arborization (DA) neurons in larval *Drosophila melanogaster* epidermis (Grueber et al., 2002). The four DA neuronal subtypes show differing capacities for homotypic receptive field arrangement. Class IV and III DA neurons tile the body wall with differing degrees of receptive field overlap, while Class I and II DA neurons show no repulsive response to homotypic neurons (Grueber et al., 2003). Tiled dendritic receptive fields have also been observed in neurons innervating the epidermis in *Manduca sexta* and *Haementeria ghilanii* (Blackshaw et al., 1982; Grueber et al., 2001; Kramer and Kuwada, 1983). The paradigm of sensory neuron receptive field patterning is also present in vertebrate systems. *Danio rerio* trigeminal neurons arrange at the midline through repulsive mechanisms like those seen in *Drosophila* (Sagasti et al., 2005). In the mouse retina, multiple examples of dendritic tiling and population-based organization exist. Bipolar cell subtypes and horizontal cells tile to innervate all available photoreceptor terminals while excluding neighboring homotypic dendritic fields and can adjust their receptive field size to accommodate changes in population. (Huckfeldt et al., 2009; Reese et al., 2005). In contrast, some amacrine cells show substantial overlap of dendritic arbors with homotypic neighbors and maintain dendritic field size even when cell density is altered (Farajian et al., 2004; Keeley et al., 2020; Lee et al., 2011).

In this study, we closely examine the development and maturation of hair follicles and follicle-innervating LTMR neurons over the first three postnatal weeks in mice. We test the hypothesis that LTMRs tile their receptive fields through a process which responds to increases in neuronal population by generating smaller receptive fields. We show that follicle innervation and hair follicle maturation coincide during the early postnatal window (before postnatal day 7, P7), and that by P14, follicle-innervating LTMR receptive fields have similar numbers of mature LLEs as adult receptive fields. Finally, using genetic tools to sparsely label single follicle-innervating LTMR neurons, we show that they form receptive fields in part through homotypic competition during the early postnatal period.

## Results

### Mouse hair follicle maturation occurs during the early postnatal window

Follicle-innervating LTMR neurons are highly branched in the skin to form receptive fields of highly specialized LLEs around hair follicles. Each follicle-innervating LTMR subtype selectively innervates specific hair follicle subtypes (Fig. 1A). However, the process by which this occurs during development is poorly understood. To understand the temporal relationship between hair follicles and LTMRs during development, we first characterized the postnatal maturation of hair follicles. Previous literature characterizing hair follicle maturation demonstrated that hair follicles are established in multiple waves starting at embryonic day 15 (E15) and continue through P9, when all hair follicles have a mature structure, including a hair shaft emerging from the skin (Paus et al., 1999). To assess hair follicle density during the early postnatal period, back skin was collected from mice at time points between P0 and P60 (Fig. 1B-G). Hair follicles could be easily identified as cell-dense structures in skin sections by staining with the nuclear marker DAPI. Hair follicle density was calculated by counting the number of hair follicle structures (long DAPI-dense structures) present in a known length of skin. Hair follicle density initially remains stable from P1-P6, then decreases beginning at P14 as mice grow more rapidly (Fig. 1H, Supplementary Table 1). This decrease in density corresponds with hair follicles completing their passage through the hair follicle cycle, as can be seen by the changes in dermal thickness over time (Müller-Röver et al., 2001).

**Figure 1.**
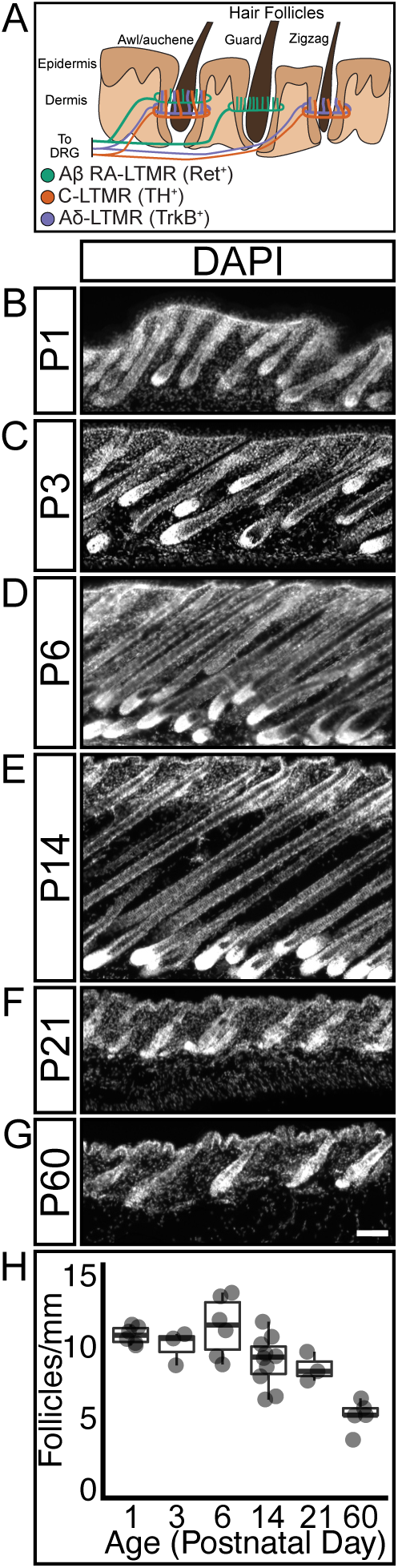
Mouse hair follicle density decreases as mice age. A) Schematic showing hair follicle-innervating LTMR neuron subtypes projecting to the skin, where their axons branch and selectively innervate specific hair follicle types. B-G) Sagittal sections of mouse back skin stained with DAPI to highlight hair follicle structures. Section height increases, then decreases as mice age and progress through the hair follicle life cycle. Scale bar – 50 μm H) Hair follicle density remains stable for the first week of postnatal life, then decreases as mice grow. N= 6(P1), 3(P3), 6(P6), 10(P14), 3(P21), 5(P60)

We further analyzed the changes in hair follicle density by normalizing the density measurements for each timepoint to the average body length of a cohort of age-matched mice (Fig. S1A). This normalization supports the conclusion that hair follicle density decreases as mice grow, and that the density of hair follicles decreases more than would be expected if mice were adding hair follicles at a rate that kept pace with animal growth (Fig. S1B). From these data, we conclude that hair follicle maturation occurs rapidly in the early postnatal window and that the rate of hair follicle addition decreases in maturing mice.

### Sensory neurons form specialized endings concurrently with hair follicle maturation

We next examined the developmental time course of hair follicle innervation using the pan-neuronal marker β-III Tubulin. We quantified the proportion of hair follicles with a β-III Tubulin-positive nerve ending encircling the follicle at multiple developmental timepoints from P1-P60 (Fig. 2A-F). Axons are present in the dermis and epidermis at P1, but very few innervate hair follicles at this age (Fig. 2A, 2G). While axon branches are present at the expected depth for follicle-innervating LLEs (inset region, Fig. 2A), these neurite branches do not yet encircle the hair follicle. There is a significant increase in the proportion of innervated hair follicles at P3, with over half of hair follicles encircled by a β-III Tubulin-positive axon (Fig. 2B, 2G). At P3 the encircling endings have not formed the LLE projections characteristic of follicle-innervating LTMRs. By P6 the innervation of hair follicles is nearly complete and the characteristic LLEs have appeared, projecting towards the epidermal surface along the hair shaft from the encircling neurite (Fig. 2C and inset, 2G). By P14, hair follicle innervation is complete, and the proportion of innervated hair follicles and LLE morphology remains unchanged at P21 and P60 (Fig. 2D-F, 2G, Supplementary Table 1). These findings are largely in agreement with other studies and suggest that murine hair follicle innervation occurs during the early postnatal window, and that by the end of the first postnatal week hair follicle innervation has reached a grossly mature state (Meltzer et al., 2022b preprint; Peters et al., 2002).

**Figure 2.**
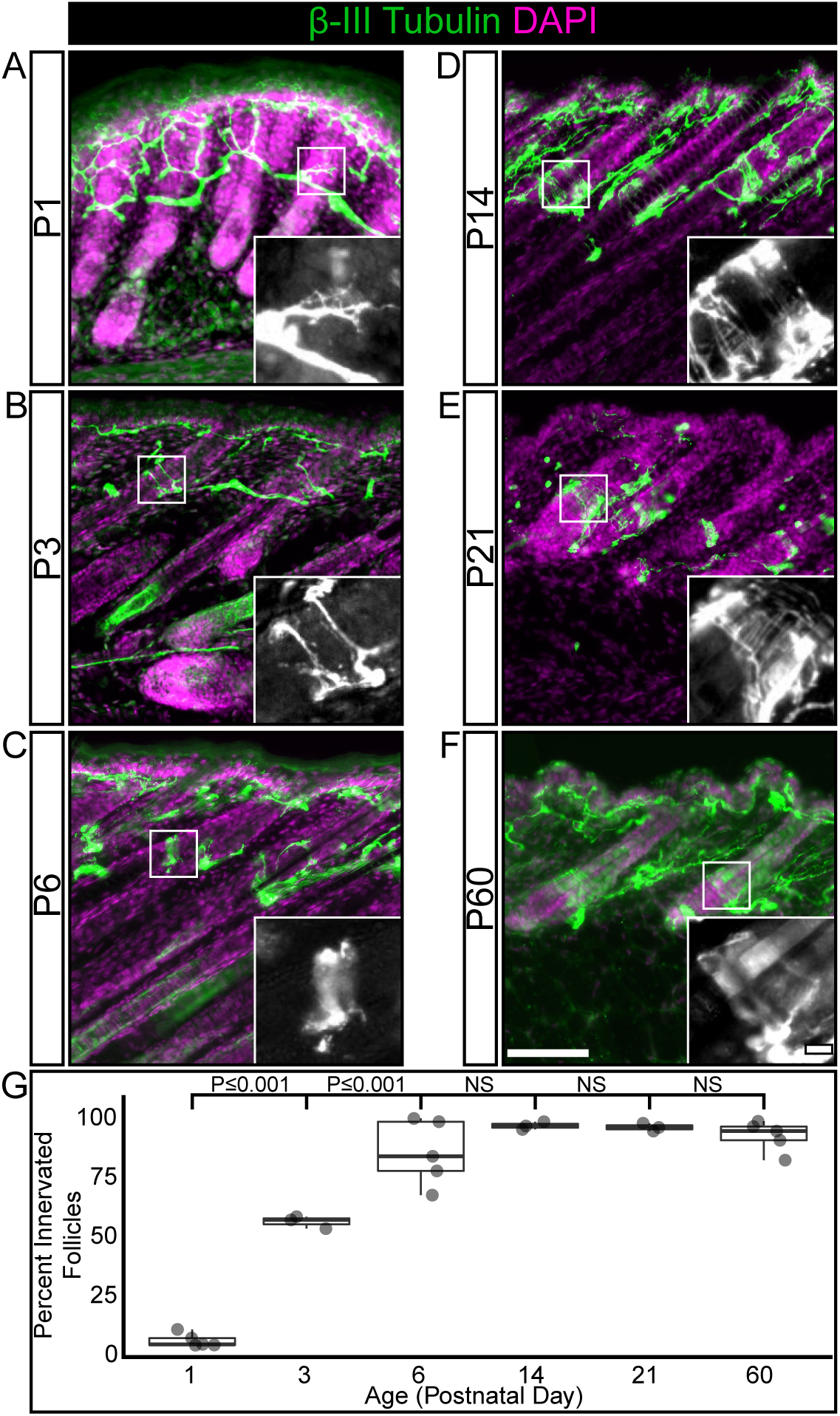
The percentage of innervated hair follicles increases during the first postnatal week, then remains stable into adulthood. A-F) Representative images of hair follicle innervation labeled with a pan-neuronal marker (β-III Tubulin, green) and DAPI (magenta). Mature longitudinal-lanceolate endings begin to appear by P6. Insets – Single channel images of follicle innervation. Scale bar – 50 μm, insets – 10 μm. G) Quantification of the percentage of hair follicles innervated by a β-III Tubulin positive axon. One-way ANOVA with Tukey’s HSD post-hoc testing (ANOVA P≤ 0.001, P1-P3 P≤ 0.001, P3- P6 P≤ 0.001) N= 5(P1), 3(P3), 5(P6), 3(P14), 3(P21), 5(P60)

### Aβ RA-LTMR follicle innervation precedes C-LTMR and Aδ-LTMR follicle innervation

We next investigated whether there is a temporal hierarchy of hair follicle innervation between LTMR subtypes. To assess the development of hair follicle innervation by specific LTMR subtypes, we utilized three separate labeling strategies. Based on data from single-cell RNA sequencing performed on adult dorsal root ganglia neurons and previous studies examining LTMR development, we identified ubiquitous markers for Aβ RA-LTMRs (anti-Calbindin immunohistochemistry) and C-LTMRs (*Vglut3^iCre^* driven *TdTomato* expression (*Ai9*)), and an inducible marker for Aδ-LTMRs (*TrkB^CreERT2^* driven placental alkaline phosphatase (*R26^iAP^*) reporter) (Badea et al., 2003; Lou et al., 2013; Rutlin et al., 2014; Seal et al., 2009; Sharma et al., 2019; Usoskin et al., 2014).

Calbindin^+^ Aβ RA-LTMRs innervate guard (tylotrich) hairs, which make up approximately 1% of the hair follicles in mouse skin, and awl/auchene hairs which make up approximately 25% of the follicles (Kuehn et al., 2019; Li and Ginty, 2014). At P1, there are very few Calbindin^+^ follicle-encircling nerve endings present (Fig. 3A, 3G), but by P3 the proportion of hair follicles with Calbindin^+^ encircling LTMR endings has significantly increased to approximately adult levels (Fig. 3B, 3G) and remains stable through at least P60 (Fig. 3C-F, 3G, Supplementary Table 1). C-LTMRs innervate the two families of non-tylotrich hairs, the zigzag hairs (∼74% of hair follicles) and the awl/auchene hairs (Kuehn et al., 2019; Li and Ginty, 2014). At P1, Vglut3^+^ C-LTMR axons are present in the skin but there are few follicle-innervating nerve endings present (Fig. 3H, 3N). The proportion of C-LTMR-innervated hair follicles significantly increases between P1 and P3, with over half of hair follicles receiving innervation from a C-LTMR axon ending (Fig. 3I, 3N). By P6 the proportion of C-LTMR innervated hair follicles has increased to approximately adult levels, where it remains stable (Fig. 3J-M, 3N, Supplementary Table 1).

**Figure 3.**
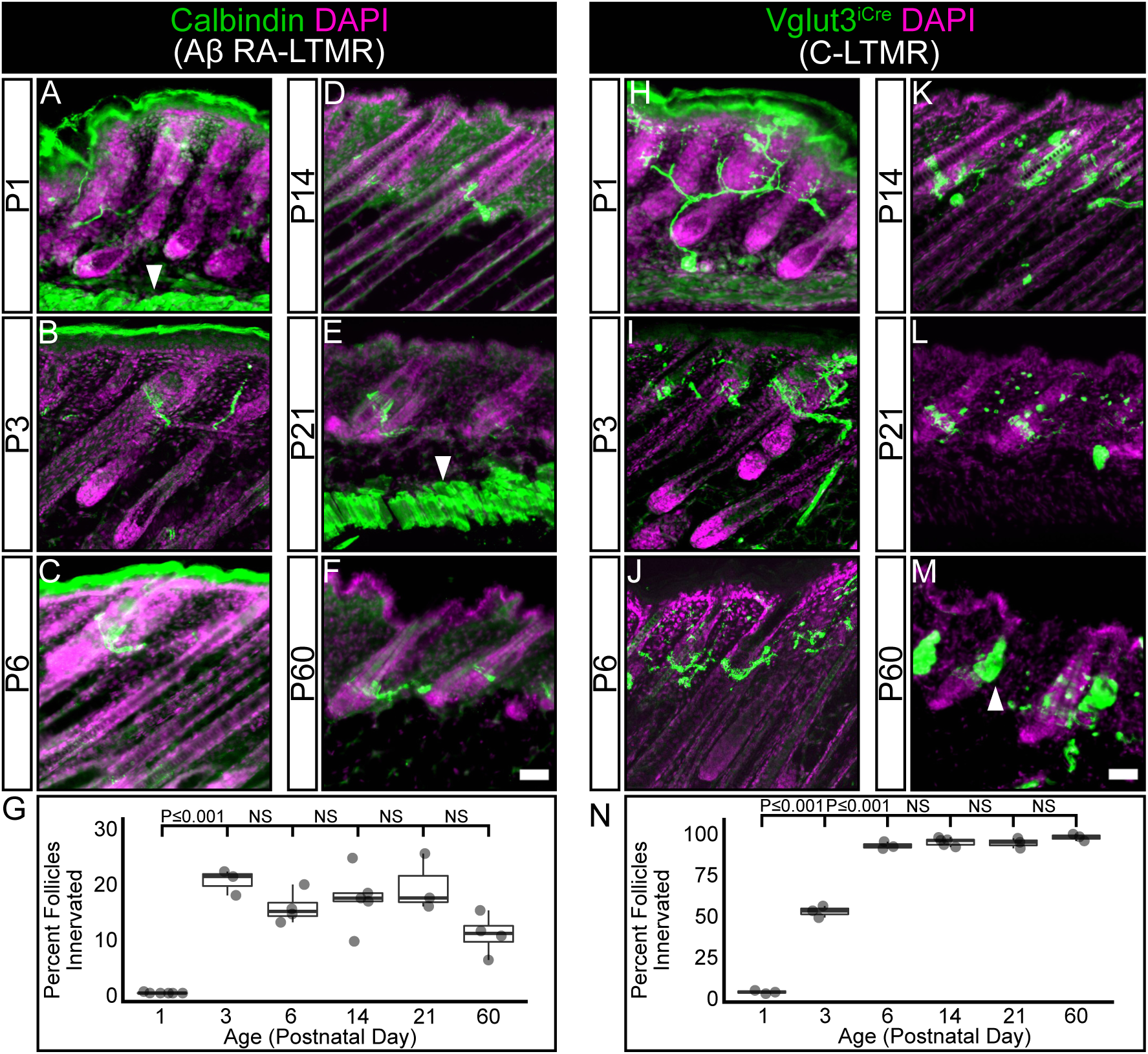
Aβ RA-LTMR neurons complete innervation of hair follicles before C-LTMR neurons. A-F) Representative images of hair follicle innervation labeled with a Aβ RA-LTMR specific marker (Calbindin, green) and DAPI (magenta). Mature longitudinal-lanceolate endings begin to appear by P3. Arrowheads (A, E) – Subcutaneous muscle autofluorescence, Scale bar – 50 μm G) Quantification of the percentage of hair follicles innervated by a Calbindin positive Aβ RA- LTMR axon. The percentage of innervated hair follicles reaches its maximum level by P3. One- way ANOVA with Tukey’s HSD post-hoc testing (ANOVA P≤ 0.001, P1-P3 P≤ 0.001). N= 6(P1), 3(P3), 4(P6), 5(P14), 3(P21), 4(P60) H-M) Representative images of hair follicles innervated by Vglut3+ axons (*Vglut3^iCre^;TdTomato*, green) and DAPI (magenta). Mature longitudinal-lanceolate endings begin to appear by P3. Arrowhead (M) – Sebaceous gland autofluorescence, Scale bar – 50 μm N) Quantification of the percentage of hair follicles innervated by a *Vglut3^iCre^;TdTomato* positive C-LTMR axon. The percentage of innervated hair follicles increases between P1 and P3, and reaches its maximum level by P6. One-way ANOVA with Tukey’s HSD post-hoc testing (ANOVA P≤ 0.001, P1-P3 P≤ 0.001, P3-P6 P≤ 0.001). N= 3(P1), 3(P3), 3(P6), 5(P14), 3(P21), 3(P60)

To explore the timing of Aδ-LTMR receptive field development, we crossed the inducible *TrkB^CreERT2^* line with a *Cre*-inducible GFP reporter line (*Ai140D*) to sparsely label Aδ-LTMRs, and imaged receptive fields at multiple time points (Daigle et al., 2018; Pomaville and Wright, 2021; Rutlin et al., 2014). The Aδ-LTMRs were ideal for developmental receptive field analysis as they were genetically accessible with prenatal tamoxifen administration and *TrkB^CreERT2^* labels a stable population of cells throughout development. In the first 24 hours after birth, there are few follicle-innervating nerve endings present in the skin, though TrkB^+^ Aδ-LTMR axons are present and have begun to branch (Fig. 4A, 4E). At P3 the neurons have begun to form follicle-innervating endings (Fig. 4B, arrowheads, 4E), but still contain many non-follicle-innervating neurites (Fig. 4B, 4E). The transition from non-ending forming neurites to follicle-innervating endings continues at P6 (Fig. 4C, 4E). By P14 the receptive fields resemble those in mature animals, with few non-follicle-innervating neurites present, and hemicircular LLEs are present at most hair follicles in the innervated area (Fig. 4D, 4E, Supplementary Table 2). The completion of Aδ-LTMR development is further confirmed in sections of P14 skin co-labeled with β-III Tubulin and *TrkB^CreERT2^;R26^iAP^*driven with multiple high prenatal doses of tamoxifen (Fig. S2A- C). Quantification of the proportion of hair follicles innervated by a TrkB^+^/β-III Tubulin ^+^ ending shows innervation matching expected proportions in adult animals (92.2% ± 3% s.e.m.). Taken together, these data show that innervation of hair follicles by LTMRs is an ongoing process during the first two postnatal weeks, and there is a subtype-specific temporal hierarchy of hair follicle innervation, with Calbindin^+^ Aβ RA-LTMRs preceding both Vglut3^+^ C-LTMRs and TrkB^+^ Aδ-LTMRs.

**Figure 4.**
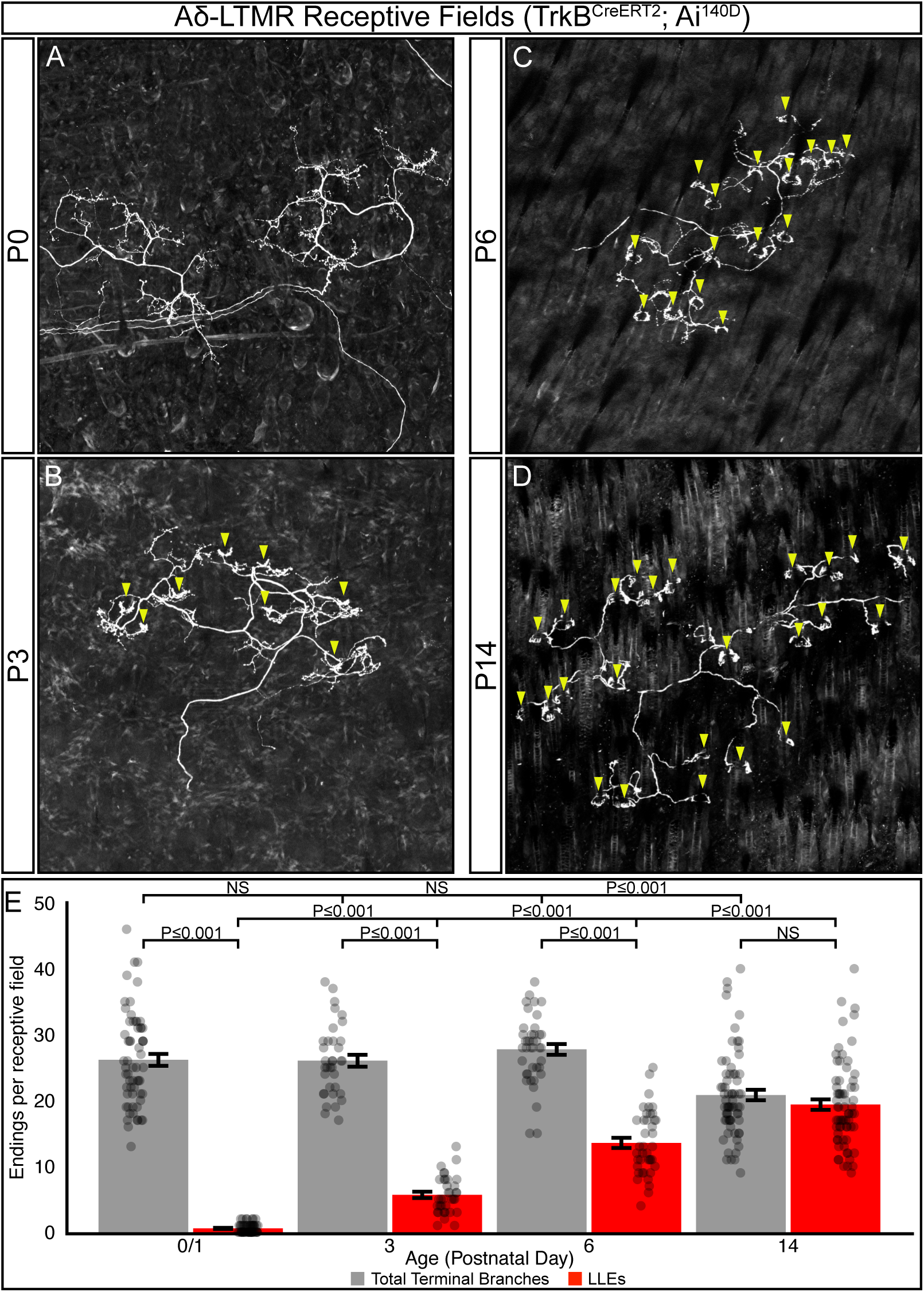
Aδ-LTMRs develop longitudinal-lanceolate endings (LLEs) after elaboration/branching in the first postnatal week. A-D) Representative images of sparsely labeled Aδ-LTMR receptive fields in mouse back skin. Aδ-LTMR axons are present in the skin at P0 (A, two distinct adjacent RFs shown). Follicle- innervating LLEs first appear by P3 (B); more follicle-innervating endings are present in P6 animals (C), and by P14 nearly all terminal branches end in LLEs (D). Arrowheads – Follicle- innervating LLEs. Scale bar – 100 μm E) Quantification of the number of terminal branches per receptive field (grey) and the number of terminal LLEs per receptive field (red). Points represent individual receptive fields. Bars represent mean ± s.e.m. Total number of receptive fields quantified and (number of animals): P0/1: 61(4), P3: 35(3), P6: 40(3), P14: 68(3) Total branches: ANOVA P≤0.001, LLEs: ANOVA P≤0.001, Total branches vs LLEs: P0/1 Wilcoxon Rank Sum Test P≤0.001, P3 Wilcoxon Rank Sum Test P≤0.001, P6 Wilcoxon Rank Sum Test P≤0.001, P14 Wilcoxon Rank Sum Test P≥0.05

### Competition between Aδ-LTMRs shapes their receptive fields

Many sensory neurons tile their receptive fields in order to reduce homotypic overlap (Grueber et al., 2001; Grueber et al., 2002; Grueber et al., 2003; Sagasti et al., 2005). Of the hair follicles innervated by C-LTMRs, <30% are innervated by multiple C-LTMRs; for Aδ-LTMRs, <10% are innervated by multiple Aδ-LTMRs (Kuehn et al., 2019). We hypothesized that this near-exclusive pattern of receptive field innervation is the result of homotypic competition for territory. If this were true, we would expect individual neurons of a given subtype to adjust their receptive field size in response to changes in neuronal subtype population. We developed a genetic strategy to test this hypothesis using deletion of the proapoptotic protein *Bax* to generate a model of neuronal overpopulation. Deletion of *Bax* blocks developmental programmed cell death in DRG neurons and results in 50-70% increases in the number of TrkA, TrkC, TrpV1, and CGRP-positive DRG neurons (Kinugasa et al., 2006; Sun et al., 2004; Suzuki et al., 2010). However, the effect of *Bax* deletion on LTMR populations has not previously been assessed.

To quantify the effect of *Bax* deletion on Aδ-LTMR and C-LTMR populations, we collected cryosections from the sixth thoracic DRG, which contains the neurons responsible for innervating the thoracic skin but contains no limb or glabrous skin innervating neurons (Takahashi and Nakajima, 1996). Sections were labeled with markers for sensory neurons (Islet 1/2, Fig. 5A, 5E), C-LTMRs (Tyrosine Hydroxylase (TH), Fig. 5B, 5F), and Aδ-LTMRs (*TrkB^CreERT2^; R26^iAP^*, (see methods) Fig. 5C, 5G), and the number of marker-positive cell bodies in the imaged sections was manually counted. Cell counts from wild-type and heterozygous mice showed no significant differences in any comparison and were pooled together for analysis (Fig. S3A-C). We found an overall 19% increase in the number of Islet 1/2-positive neurons in *Bax^-/-^* animals compared to control animals. The number of *TrkB^CreERT2^*^+^ Aδ-LTMRs showed a significant increase (53%) in *Bax^-/-^* animals compared to control littermates. The number of TH^+^ C-LTMRs increased 33% in *Bax^-/-^* animals compared to control littermates, which was not statistically significant (Fig. 5I, Supplementary Table 3). Deletion of *Bax* had no effect on hair follicle density in P21 mice, as measured using Oil Red O staining (Fig. S4A-D). Therefore, *Bax* deletion results in a significant overpopulation of Aδ-LTMR neurons but does not significantly affect the number of C-LTMR neurons or the targets of LTMR innervation (hair follicles).

**Figure 5.**
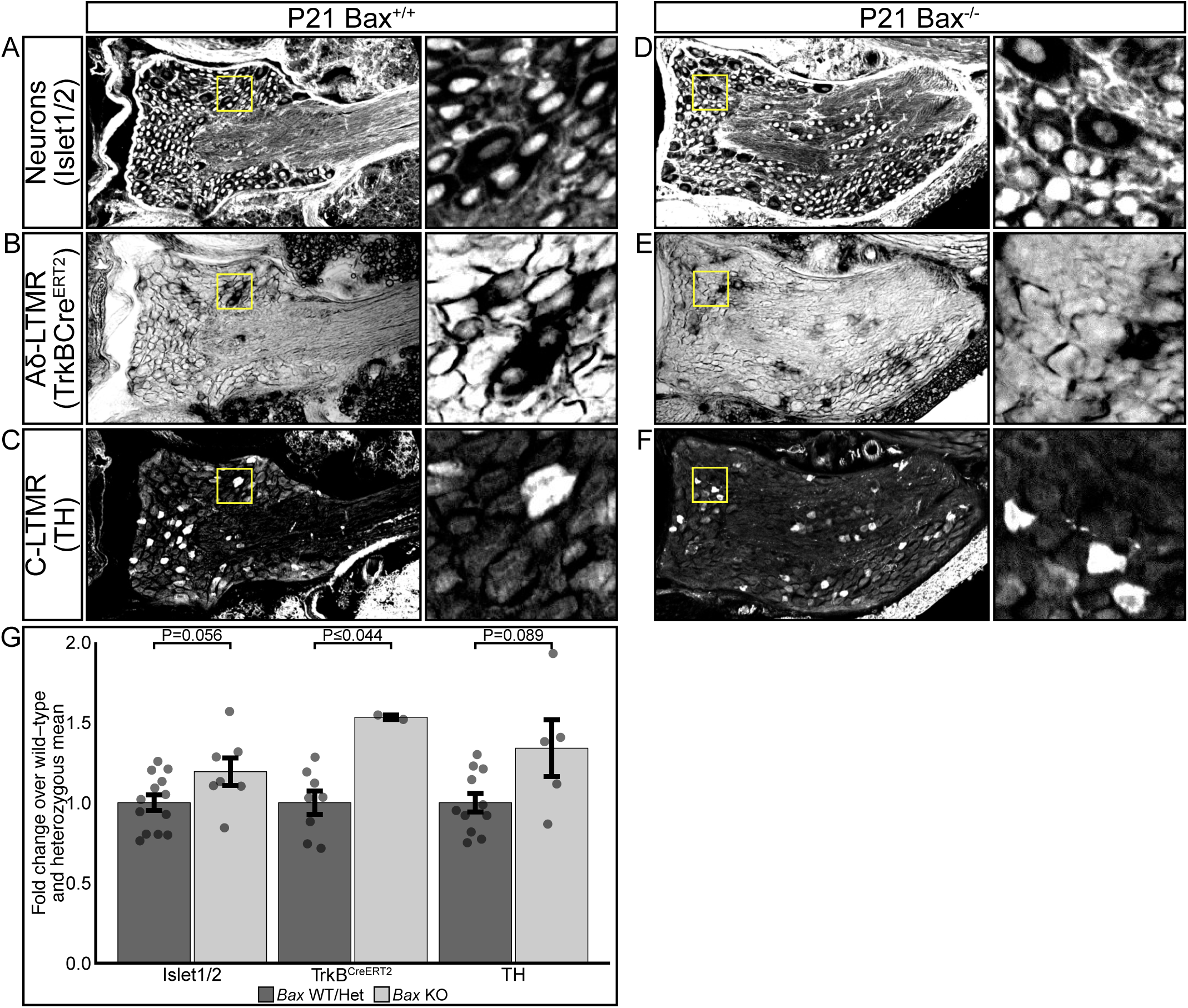
Genetic deletion of the proapoptotic protein *Bax* has differential effects on LTMR populations. A-F) Representative sections of T6 dorsal root ganglia from 21-day old wild-type littermates (A- C) and *Bax* knockout (D-F) mice labeled with markers for neuronal nuclei (A, D Islet 1/2), Aδ- LTMR cell bodies (B, E TrkB^CreERT2^; R26^iAP^), C-LTMR cell bodies (C, F Tyrosine Hydroxylase). Scale bar – 50 μm. Insets show high-magnification of boxed regions. E) Quantification of the change in T6 DRG LTMR neuron proportions in *Bax* knockout animals over wild-type and heterozygous animals. Bars represent mean ± s.e.m. N = Islet 1/2: WT/Het – 13, KO – 7, TrkB: WT/Het – 8, KO – 2, TH: WT/Het – 11, KO – 5. Islet 1/2 Wilcoxon Rank Sum: P=0.056, TrkB^CreERT2^ Wilcoxon Rank Sum: P≤0.044, TH Wilcoxon Rank Sum: P=0.089

If LTMRs of the same subtype compete for territory, we would expect that the receptive fields of individual Aδ-LTMRs would become smaller in *Bax^-/-^* animals to accommodate the excess number of neurons, while the receptive fields of C-LTMRs would remain largely unchanged. To test this, we generated triple transgenic mice carrying 1: tamoxifen inducible *Cre* driver lines for Aδ-LTMRs (*TrkB^CreERT2^*) or C-LTMRs (*TH^IRES-CreER^*), 2: a *Cre-*dependent placental alkaline phosphatase reporter (*R26^iAP^*), and 3: *Bax^+/+^, Bax^+/-^*, or *Bax^-/-^* (Badea et al., 2009; Knudson et al., 1995; Rotolo et al., 2008; Rutlin et al., 2014). We administered low doses of tamoxifen to timed-pregnant dams (Aδ-LTMRs) or juvenile mice (C-LTMRs) to sparsely label receptive fields and quantified the number of hair follicles in each receptive field of young adult animals (P21). Representative images of Aδ-LTMR and C-LTMR receptive fields (Fig. 6A-6G) from *Bax^+/+^, Bax^+/-^*, and *Bax^-/-^* mice were traced using the FIJI plugin Simple Neurite Tracer (Fig. 6A’-G’) to better illustrate the receptive field structure (Longair et al., 2011; Schindelin et al., 2012). Aδ-LTMRs showed a significant reduction in the number of hair follicles innervated by each neuron in *Bax^-/-^* animals (24.2 ± 0.5 s.e.m. follicles/neuron) compared with both wild-type (31.6 ± 0.8 s.e.m. follicles/neuron) and heterozygous (29.5 ± 0.5 s.e.m. follicles/neuron) animals (Fig. 6D). This represents a 23% decrease in the number of hair follicles innervated by single Aδ-LTMRs in *Bax^-/-^* animals. In contrast to the Aδ-LTMRs, there was no significant difference in the number of hair follicles innervated by each C-LTMR receptive field in *Bax^-/-^* (14.8 ± 0.7 s.e.m. follicles/neuron), heterozygous (16.6 ± 1.1 s.e.m. follicles/neuron), or wild-type (16.6 ± 0.5 s.e.m. follicles/neuron) animals (Fig. 6H). No significant differences were seen between mice of the same *Bax* genotype across litters for both Aδ-LTMRs and C-LTMRs (Fig. S5A-B).

**Figure 6.**
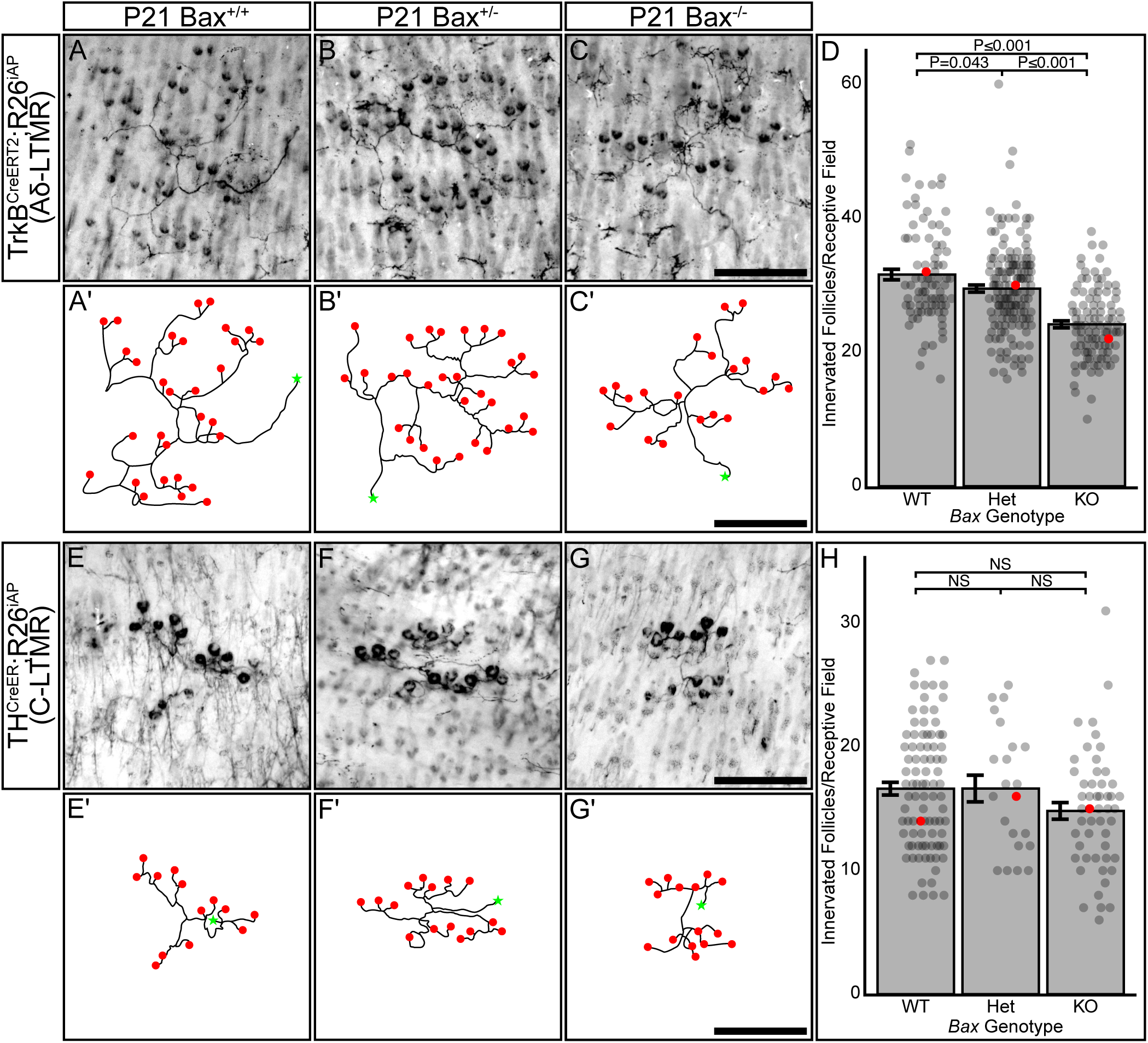
Aδ-LTMRs adjust their receptive field size to accommodate changes in neuronal population due to *Bax* deletion. A-C’) Representative images and traced reconstructions of isolated Aδ-LTMR receptive fields from back skin of P21 *TrkB^CreERT2^; R26^iAP^*; *Bax* wild-type (A, A’), heterozygous (B, B’), and knockout (C, C’) animals. Red dots in traces correspond with hair follicles, green stars indicate the axon segment from which the traced RF originates. Scale bar – 250 μm D) Quantification of the number of hair follicles innervated by each Aδ-LTMR receptive field in *TrkB^CreeRT2^;R26^iAP^*; *Bax* wild-type, heterozygous, and knockout animals. Receptive fields in *Bax* knockout animals innervate fewer hair follicles than wild-type and heterozygous animals. Each dot represents one receptive field, red dots represent the data point from to the associated representative image. Bars represent mean ± s.e.m. One-way AONVA with Tukey’s HSD post- hoc testing ANOVA P≤0.001, WT-KO P≤ 0.001, Het-KO P≤ 0.001, WT-Het P= 0.043. Number of individual receptive fields (number of animals): WT: 85(3), Het: 154(5), KO: 103(3) E-G’) Representative images and traced reconstructions of isolated C-LTMR receptive fields from back skin of P21 *TH^CreERT2^; R26^iAP^*;*Bax* wild-type (E, E’), heterozygous (F, F’), and knockout (G, G’) animals. Red dots in traces correspond with hair follicles, green stars indicate the axon segment from which the traced RF originates. Scale bar – 250 μm H) Quantification of the number of hair follicles innervated by each C-LTMR receptive field in *TH^CreERT2^; R26^iAP^*;*Bax* wild-type, heterozygous, and knockout animals. No significant difference was seen in the number of hair follicles innervated across *Bax* genotypes. Each dot represents one receptive field, the red dots represent the data point from to the associated representative image. Bars represent mean ± s.e.m. One-way AONVA, P= 0.098. Number of individual receptive fields (number of animals): WT: 90(6), Het: 22(4), KO: 51(3)

Therefore, we conclude that follicle-innervating LTMRs form their receptive fields in part through homotypic population-based competition.

## Discussion

The organized development of skin- and hair follicle-innervating sensory neurons is essential for communicating meaningful information about touch to the brain in order to moderate behavior. In this study we make use of multiple cell-type specific molecular and genetic tools to analyze the development of hair follicle innervation and explore the principles that govern follicle-innervating LTMR receptive field organization. We show innervation of hair follicles and initial LLE development occurs during the first two postnatal weeks, and that by P14, mice show an adult innervation pattern. We also show that LTMR subtypes have a differential dependence on Bax-mediated developmental apoptosis, with Aδ-LTMRs responding to deletion of *Bax* by significantly increasing their population, while the population of C-LTMRs increases only slightly. Finally, we demonstrate that in response to this change in population, Aδ-LTMR neurons shrink their receptive fields to accommodate their neighbors.

### LTMR receptive fields undergo gradual refinement during the first two postnatal weeks

Previous work has described the tight association between hair follicles and the follicle-innervating LTMRs, showing PGP9.5 labeled axons wrapping hair follicles as early as E18, and describing the adult morphology of LTMRs in a population-specific manner (Bai et al., 2015; Li and Ginty, 2014; Li et al., 2011; Peters et al., 2002; Rutlin et al., 2014). Our work builds upon this by analyzing the development of follicle innervation during the first three postnatal weeks in an LTMR-subtype-specific manner. Specifically, we present a detailed timeline showing a gradual development of follicle innervation by Aβ RA-LTMRs and C-LTMRs during the first two postnatal weeks (Fig. 3). We also used a genetically-driven sparse labeling approach to examine the development of individual Aδ-LTMR receptive fields, showing a gradual development of follicle-innervating endings and refinement of LTMR receptive field characteristics during the first two postnatal weeks (Fig. 4). These findings agree with recent work describing interactions between LTMR axons and terminal Schwann cells at the hair follicle during this time period (Meltzer et al., 2022b preprint).

### Follicle Innervation Develops in a Temporally Ordered Manner

Our detailed examination of LTMR follicle innervation development revealed a temporal hierarchy for the different subtypes, with Aβ RA-LTMRs completing their innervation of hair follicles by P3, before C-LTMRs and Aδ-LTMRs do at P6 (Fig. 3). Recent comparison of the development of Aβ RA-LTMRs and Aδ-LTMRs supports this conclusion, showing Aβ RA-LTMRs maturing at P3 and Aδ-LTMRs maturing after P6 (Meltzer et al., 2022b preprint). This temporal hierarchy parallels the maturation of hair follicle subtypes. Guard hair placodes, which are innervated by Aβ RA-LTMRs, consolidate between E14 and E15, with the first guard hair shafts appearing between E16 and E18. The consolidation of other hair follicle placodes, which are innervated by C-LTMRs and Aδ-LTMRs, occurs in successive waves continuing until after birth (Hardy, 1949; Mann, 1962; Peters et al., 2002). This temporal hierarchy of innervation could also be due to the temporal order of LTMR differentiation. EdU birth dating studies have shown that Aβ RA-LTMR neurons are born as early as E9.5 and have a peak birth rate at E10.5 before sharply dropping off. In contrast, C-LTMRs and Aδ-LTMRs are born approximately one day later, peaking at E11.5 and gradually tapering off by E13.5 (Landy et al., 2021). Furthermore, single-cell transcriptomic analysis of the developing DRG shows that C-LTMRs are among the last populations to mature as transcriptionally distinct subtypes, mirroring their birthdate hierarchy (Sharma et al., 2019). It is tempting to speculate that early-born Aβ RA-LTMRs could be a “pioneer” axon population in the periphery, functioning as a scaffold along which axons from Aδ-LTMRs and C-LTMRs grow before branching to form receptive fields. In invertebrate systems, pioneer axons play a critical role in establishing peripheral innervation patterns, and ablation of these cells leads to significant disruptions in follower axons (Bate, 1976; Edwards, 1977; Edwards et al., 1981; Keshishian, 1980; Klose and Bentley, 1989). In vertebrates, early extending axons often act as a permissive substrate upon which later-born neurons can extend their axons, but the necessity of these early-extending axons is less clear (Melançon et al., 1997; Pike et al., 1992; Pittman et al., 2008). In an interesting parallel, Ret is expressed in both early-born Aβ RA-LTMRs and in a population of pioneer axons in the zebrafish posterior lateral line (Tuttle et al., 2019). Future studies will be required to determine if early born Ret^+^ Aβ RA-LTMRs function as pioneers in the mouse DRG.

### Knockout of the proapoptotic protein *Bax* shows differential effects on LTMR populations

To understand how competition between LTMR neurons might influence receptive field development, we devised a genetic strategy to drive LTMR overpopulation by blocking developmental apoptosis. Multiple studies have demonstrated that sensory neurons are initially overproduced, followed by a wave of developmental apoptosis regulated by target-derived factors which ensures neurons and targets are appropriately matched (Reviewed in Buss et al., 2006; Fariñas et al., 1994; Oppenheim et al., 1991). Genetic deletion of the pro-apoptotic protein *Bax* blocks developmental apoptosis and results in significantly increased numbers of neurons in the DRG (Patel et al., 2000; White et al., 1998). While *Bax* knockout animals show a 1.5-fold to 1.8-fold increase in the population of proprioceptive and nociceptive DRG neurons and increased sensory axon density in the paw, previous studies had not examined the effect of *Bax* deletion on populations of LTMRs (Suzuki et al., 2010). Our experiments specifically examining follicle-innervating LTMRs showed a significant increase in the number of Aδ-LTMR (*TrkB^+^*) DRG neurons, but only a small, non-significant increase in C-LTMR (*TH^+^*) DRG neurons. These results reveal an interesting difference in the dependence of different neuronal LTMR subtypes on developmental apoptosis to achieve their final numbers. The basis for this difference is unknown, but may involve mechanisms related to the specification, differentiation, and/or organization of LTMR neurons, such as the role for target-derived trophic cues in LTMR population determination and LLE maintenance.

### Homotypic competition plays a role in LTMR receptive field organization

The development of tools for subtype-specific labeling of LTMRs revealed several interesting organizational properties. First, different LTMR subtypes have different hair follicle innervation patterns, with Aβ RA-LTMRs innervating guard and awl/auchene hairs, while Aδ- LTMRs and C-LTMRs innervate zigzag and awl/auchene hairs. Second, LTMRs have distinct, highly stereotyped receptive fields that typically innervate a fixed range of hair follicles (Bai et al., 2015; Li and Ginty, 2014; Li et al., 2011; Rutlin et al., 2014). In our study, we attempted to address these properties at a cellular level by examining the role of homotypic competition in the establishment of receptive fields. We reasoned that inducing overpopulation of LTMR subtypes with *Bax* deletion would lead to two potential outcomes. Either neurons would decrease their receptive field size to accommodate the increased number of neighbors and preserve homotypic tiling, or they would maintain their receptive field size at the expense of tiling. Aδ-LTMRs showed a clear preference for adjusting their receptive field size to accommodate the population increase, with each neuron in *Bax^-/-^* mice forming a receptive field innervating significantly fewer hair follicles than in wild-type mice. In contrast, despite a small increase in the number of C-LTMRs in *Bax* deficient mice, these neurons did not adjust their receptive field size, suggesting they may overlap more than in *Bax* wild-type animals. This difference in response could be due to the stringency of the tiling behaviors of the two LTMR subtypes: approximately 10% of hair follicles in wild-type mice are innervated by two distinct Aδ- LTMRs, whereas 25% of hair follicles are innervated by two distinct C-LTMRs (Kuehn et al., 2019). We attempted a complementary approach, ablating a portion of the Aδ-LTMR population prenatally using the *TrkB^CreERT2^*driver crossed to a *Cre*-dependent sensory neuron specific Diphtheria toxin receptor (*Avil^iDTR^*), reasoning that population depletion before follicle innervation would result in larger receptive fields (Stantcheva et al., 2016). Unfortunately, these experiments resulted in perinatal lethality, potentially due to off-target effects of Diphtheria toxin administration.

At the molecular level, there is little known about the cues that regulate the targeting of LTMR subtypes to specific hair follicle types or the establishment of receptive fields. While BDNF derived from hair follicle-associated epithelial cells is important for the polarized innervation pattern and LLE maturation of TrkB^+^ Aδ-LTMR axonal endings, it is dispensable for the initial attraction of Aδ-LTMR axons to hair follicles, and it is unknown whether it regulates receptive field size (Rutlin et al., 2014). Similarly, recent work has shown that the GPI-linked receptor Netrin-G1 is required in LTMRs for the proper maturation of terminal endings around hair follicles, but not for terminal axon branching or receptive field size (Meltzer et al., 2022b preprint). In contrast, ɣ-Protocadherins are important for the peripheral branching and innervation of non-guard hair follicles by Aβ-LTMRs, but not for the maturation of LLEs (Meltzer et al., 2022a preprint). Together, these data suggest that the initial innervation of hair follicles and the morphological maturation of their terminal endings are distinct events regulated by different molecular pathways. Interestingly, terminal Schwann cells are the source for ligands/binding partners for both Netrin-G1 and ɣ-Protocadherins in LTMRs, demonstrating that they play a critical role in regulating hair follicle innervation (Meltzer et al., 2022a preprint; Meltzer et al., 2022b preprint).

### Why do distinct LTMR subtypes display different receptive field morphologies?

Presumably, tight regulation of LTMR receptive field size and organization is intrinsically tied to their sensory functions. In order to allow for precise localization of a stimulus, cutaneous sensory neurons must innervate a constrained area of the skin and relay this information to the proper regions of the central nervous system. LTMR receptive field size and organization is also an important factor in determining the firing properties of individual neurons. Single neuron recordings from LTMRs have demonstrated significant sensory specialization across subtypes. The Aβ RA-LTMRs and C-LTMRs both fire action potentials in response to generalized skin indentation and skin stroking in the direction of hair growth (head to tail), differing in their specific response properties (Li et al., 2011). Aδ-LTMRs also fire action potentials in response to skin indentation, but selectively fire in response to skin stroking opposite the direction of hair growth (tail to head) due to their hemicircular arrangement on the caudal side of the hair follicle (Rutlin et al., 2014). Aβ RA-LTMRs, Aδ-LTMRs, and C-LTMRs can all fire action potentials through the stimulation of a single hair follicle. In contrast, the Aβ Field-LTMRs, which have significantly larger receptive fields than other LTMR subtypes, can only fire action potentials through summation when multiple hairs are stimulated simultaneously (Bai et al., 2015). These unique properties result in a sensory system able to accurately localize higher magnitude stimuli through small receptive fields, while also detecting stimuli too faint to trigger action potentials through a single LLE.

The final factor potentially involved in LTMR tiling is an inherent characteristic of the LTMR receptive fields themselves. In adult mice, LTMR axons branch and form highly stereotyped receptive fields, with some subtypes such as the C-LTMRs forming fields innervating on average 15 hair follicles, while on the other extreme the Aβ Field-LTMRs can innervate over 150 hair follicles (Bai et al., 2015). Work examining the tiling of LTMR receptive fields has shown an interesting correlation between the number of hair follicles each individual receptive field innervates and the degree of overlap between neighboring homotypic receptive fields. Aβ Field-LTMRs, which have the largest receptive fields, show the least degree of overlap with homotypic neurons. C-LTMRs, which have the smallest receptive fields, show the highest degree of overlap (Kuehn et al., 2019).

The ordered development of follicle innervation and interactions between neighboring homotypic neurons during receptive field development raises the question of whether similar mechanisms regulate the organization of central DRG afferents in the spinal cord dorsal horn. The different LTMR subtypes target their axons to specific lamina in the dorsal spinal cord, somatotopically arrange their axons, and show a high degree of synaptic partner specificity (Abraira et al., 2017; Li et al., 2011; Odagaki et al., 2018). The cellular and molecular mechanisms that govern central afferent organization remain largely unknown, but recent advancements in imaging and genetic tools may open this field to further investigation.

## Methods

### Mouse Husbandry

Mice (*Mus musculus*) were housed in standard conditions, cared for by the Department of Comparative Medicine at Oregon Health and Science University. All animal experimental procedures were approved by OHSU Institutional Animal Care and Use Committee (Protocol # IS00000539) and adhered to the NIH *Guide for the care and use of laboratory animals*. Mice were maintained on a 12-hour light/dark cycle and were provided food and water *ad libitum*. For embryonic tamoxifen administration, the morning of vaginal plug observation was designated as E0.5. Mice were maintained on a mixed genetic background due to experiments being conducted with compound transgenic lines. Mice of both sexes were used, and littermates were used as controls.

Due to the estrogen inhibiting effects of tamoxifen during pregnancy, many female mice fail to initiate labor at the proper time resulting in fetal demise. For all female animals given tamoxifen during pregnancy, pups were delivered by cesarian section at embryonic day 19.5 after mild isoflurane-induced anesthesia and rapid cervical dislocation. After removal from the uterus and amniotic sac, pups were dried gently with a paper towel and warmed on a heating pad at 37° C. Viable pups breathe spontaneously within 1-2 minutes of removal from the amniotic sac and respond to gentle tactile stimuli within 2-3 minutes, becoming pink and actively moving within 5- 10 minutes. Viable pups were placed with the litter of a nursing female mouse approximately 15 minutes after delivery and an equal number of pups were removed from the foster dam’s litter. The day of birth or cross-fostering of animals was designated as P0.

### Mouse Lines

Mouse lines used in this study have all been previously described (Table 1).

**Table 1.** Mouse Lines Used:

| Common name | Strain name | Reference | Jax/MGI Reference number |
| --- | --- | --- | --- |
| TH<sup>CreER</sup> | B6;129-*Th*<sup>tm1(cre/Esr1)Nat</sup>/J | (Rotolo et al., 2008) | JAX: 008532 |
| TrkB<sup>CreERT2</sup> | B6.129S6(Cg)-<br>*Ntrk2*<sup>tm3.1(cre/ERT2)Ddg</sup>/J | (Rutlin et al., 2014) | JAX: 027214 |
| Ret<sup>CreERT2</sup> | Ret<sup>tm2(cre/ERT2)Ddg</sup> | (Luo et al., 2009) | MGI: 4437245 |
| Vglut3<sup>iCre</sup> | Tg(Slc17a8-icre)1Edw/SealJ | (Grimes et al., 2011) | JAX: 018147 |
| Bax KO | B6.129X1-Bax<sup>tm1Sjk</sup>/J | (Knudson et al., 1995) | JAX: 002994 |
| Ai140D | B6.Cg-<sup>lgs7tm140.1(tetO-EGFP,CAG-tTA2)Hze</sup>/J | (Daigle et al., 2018) | JAX: 030220 |
| R26-iAP | B6;129-Gt(ROSA)26Sor<sup>tm2Nat</sup>/J | (Badea et al., 2003) | JAX: 009253 |

### Tamoxifen Administration

Tamoxifen stock was made by dissolving tamoxifen powder in freshly opened 100% ethanol at a concentration of 150 mg/mL and stored at -80° C until needed. For tamoxifen administration to pregnant female mice, progesterone and β-estradiol were co-administered at a ratio of 1000:500:1, Tamoxifen:progesterone:β-estradiol. Administration timing and dosage are given in associated figure legends. For all administration methods and times, the Tamoxifen/ethanol stock solution was dissolved in sunflower seed oil by vortexing until an emulsion formed. The oil/ethanol mixture was then centrifuged in a heated vacuum centrifuge (SpeedVac) for 15 minutes to evaporate residual ethanol. Pregnant female mice were administered tamoxifen through oral gavage of 100 μL of sunflower seed oil with dissolved tamoxifen-progesterone-β- estradiol mixture. P12-16 pups were administered tamoxifen through oral gavage of 50 μL of sunflower seed oil with dissolved tamoxifen. P0-P11 pups were administered tamoxifen through IP injection of 10 μL of sunflower seed oil with dissolved tamoxifen.

Tamoxifen doses and timepoints for each use case can be found in Table 2.

**Table 2.** Tamoxifen Administration Doses and Routes:

| Experiment | Labeling Density | Associated Figure | Mouse Line | Tamoxifen Dose | Admin. Route | Admin. Age |
| --- | --- | --- | --- | --- | --- | --- |
| Aδ-LTMR labeling in sections | High | Fig. S2 | TrkB<sup>CreERT2</sup>;R26<sup>iAP</sup> | 2 doses of 2 mg | P.O. (Maternal) | E12.5, E13.5 |
| Aδ-LTMR fluorescent sparse labeling | Low | Fig. 4 | TrkB<sup>CreERT2</sup>;Ai140D | 0.75 mg | P.O. (Maternal) | E13.5 |
| Aδ-LTMR Cell Body Labeling | High | Fig. 5 B, E, F | TrkB<sup>CreERT2</sup>;R26<sup>iAP</sup> | 3 doses of 0.5 mg | I.P. | P6, P7, P8 |
| Aδ-LTMR<br>RF sparse<br>labeling | Low | Fig. 6 A-D | TrkB<sup>CreERT2</sup>; R26<sup>iAP</sup> | 0.5 mg | P.O.<br>(Maternal) | E13.5 |
| C-LTMR<br>RF sparse<br>labeling | Low | Fig. 6 E-H | TH<sup>CreER</sup>; R26<sup>iAP</sup> | 2 doses of<br>2 mg | P.O. | P14, P15 |

**Table 3.** Antibodies Used:

| Antibody | Supplier | Item Number | RRID | Use concentration |
| --- | --- | --- | --- | --- |
| Rb α-Calbindin | Swant | CB 38 | AB_10000340 | 1:1000 |
| Rb α-TH | Millipore | AB152 | AB_390204 | 1:1000 |
| Sh α-TH | Millipore | AB1542 | AB_90755 | 1:500 |
| Ms α-Isl1/2 | DSHB | 39.4D5 | AB_2314683 | 1:250 |
| Gt α-TdTomato | Biorbyt | orb182397 | AB_2687917 | 1:1000 |
| Ck α-GFP | Abcam | ab13970 | AB_300798 | 1:1000 |
| Dk α-Rb IgG 488 | ThermoFisher | A-21206 | | 1:500 |
| Dk α-Rb IgG 546 | ThermoFisher | A10040 | | 1:500 |
| Dk α-Rb IgG 647 | ThermoFisher | A-31573 | | 1:500 |
| Dk α-Ms IgG 647 | ThermoFisher | A-31571 | | 1:500 |
| Dk α-Ck IgY 488 | Jackson Immuno Research | NC0215979 | | 1:500 |
| Dk α-Ck IgY Cy3 | Millipore Sigma | AP194C | | 1:500 |
| Dk α-Sh 488 | ThermoFisher | A-11015 | | 1:500 |
| Dk α-Gt 546 | ThermoFisher | A-11056 | | 1:500 |

### Tissue dissection and fixation

See (Pomaville and Wright, 2021) for a detailed description of histology methods. Briefly, animals were euthanized in accordance with IACUC approved protocols. Animals P6 and older were depilated using a commercial depilatory cream (Nair) and were washed with gentle soap and room temperature water. Skin was dissected away from the animal and placed onto a filter paper lined silicone-bottomed dish with a cover. Large fat deposits were trimmed away from the subcutaneous tissue using small scissors and the skin was pinned to the plate, subcutaneous side up, using fine insect pins. Skin was fixed 6 hours to overnight at 4° C in 4% paraformaldehyde with gentle agitation. After fixation, skin was washed with PBS for 30 minutes at room temperature with agitation before progressing to preparation for specific analysis methods. After washing the subcutaneous side of the fixed skin was scraped with a razor blade to remove remaining subcutaneous fat and muscle.

Spinal columns were dissected out and fixed overnight in 4% paraformaldehyde at 4° C with agitation. After fixation spinal columns were washed in PBS for 30 minutes at room temperature. Spinal columns were incubated overnight in 10% w/v EDTA/10% v/v glycerol in PBS to decalcify the bones for sectioning through the spinal column (DRG section immunofluorescence and AP staining). Spinal columns were washed 4-5 hours in PBS at 4° C to remove EDTA and glycerol before further processing.

### Whole Mount Skin Immunofluorescence

Fixed skin prepared as above was washed for 8 hours at room temperature in 0.2% v/v Triton X-100 in PBS with gentle agitation, changing the detergent every hour for the first 4 hours, then twice more. Skin was moved to blocking solution (5% v/v Normal Donkey Serum, 5% v/v DMSO, 0.25% v/v Triton X-100 in PBS) overnight at 4° C with gentle agitation. Skin was incubated in primary antibodies in blocking solution at the dilutions listed in Table 4 for 5 days at 4° C with gentle agitation. Skin was washed 2 hours with 0.2% v/v Triton X-100 in PBS, then 3 hours in PBS changing the PBS every hour at room temperature with agitation. Skin was moved to secondary antibodies in blocking solution at a 1:500 concentration for 2-3 days at 4° C with gentle agitation. Skin was washed 1 hour in 0.2% v/v Triton X-100, then 2-3 hours in PBS at room temperature before dehydration in a methanol gradient (30 minutes - 50% MeOH in PBS, 2 hours – 80% MeOH in PBS, overnight – 100% MeOH). After dehydration, skin was cleared in BABB and imaged.

### Whole Mount Skin Oil Red O Staining and Imaging

Fixed and washed skin samples were incubated in 60% v/v isopropanol in water for 10 minutes at room temperature with agitation. Skin was then stained in 0.3% Oil Red O (ORO) in 60% isopropanol at room temperature until dark red staining of sebaceous glands occurred (30 minutes to 3 hours). Skin was then washed twice with 60% isopropanol for 10 minutes each, then moved to water to store until imaging.

ORO stained skin was imaged by mounting epidermis side up on a glass slide with water and a standard coverslip. A 2 cm^2^ area of skin was imaged using a 5x objective and the number of hair follicles were counted in 10 randomly selected 1 mm^2^ fields. The counted field values were averaged together to generate an animal follicle density value.

### Whole Mount Skin Alkaline Phosphatase Staining

Fixed, washed, and scraped skin was heat treated in 1 mM MgCl2 in PBS at 70° C for 90 minutes with agitation every 15 minutes. Skin was allowed to cool to room temperature, then moved to a tube containing 0.1 M Tris/50 mM MgCl2/0.1 M NaCl in water, pH 9.5 with 16.6 µg/mL BCIP and 33.3 µg/mL NBT and 12.5 mM Levamisole for the AP reaction. The AP staining was allowed to develop at room temperature with agitation until labeled neurons were clearly visible in the skin. Fresh buffer and substrate were added after 24 hours if staining wasn’t satisfactory. After staining progressed to a satisfactory level, skin was pinned to silicone-bottomed dishes with insect pins and dehydrated for 30 minutes in 50% methanol in water, then 2 hours in 80% methanol in water, then overnight at room temperature in 100% methanol. Skin samples were cleared in a 2:1 mixture of Benzyl Benzoate:Benzyl Alcohol (BABB) for 15 minutes or until optically clear before mounting on a slide in BABB with a standard coverslip for imaging.

### Whole Mount Skin Imaging and Receptive Field Quantification

Receptive fields in cleared AP-stained skin were imaged using a Zeiss AxioZoom V.16 Macroscope. The number of LLEs per receptive field were manually counted using the Cell Counter plugin in FIJI/ImageJ. Receptive fields in representative images were semi-manually traced using the Simple Neurite Tracer plugin in Fiji/ImageJ (Longair et al., 2011). Receptive fields in cleared immunohistochemically labeled skin were imaged on a Zeiss AxioImager M.2 microscope with an Apotome.2 structured illumination module. The number of LLEs and non- LLE terminal branches were counted manually using the Cell Counter plugin in FIJI/ImageJ.

### Skin Section Cryosection Preparation

A 1 cm x 1 cm square of back skin from the anterior midline was taken from fixed mouse skin. The small portion was cryoprotected in a sucrose gradient, 30 minutes in 10% sucrose in PBS, 2 hours in 15% sucrose in PBS, overnight at 4° C in 20% sucrose in PBS. Samples were mounted in OCT media and oriented with the sagittal plane as the cutting face. Samples were rapidly frozen in methylbutane chilled on dry ice. 50 μm sagittal sections were cut, every fifth section mounted on a slide, and allowed to dry at room temperature 2-3 hours before moving to a freezer or processing for immunohistochemistry or AP staining as described below.

### Spinal Cord/DRG section preparation

The T5-T7 segment was dissected from fixed and decalcified spinal columns by cutting through the intervertebral disks between T4/5 and T7/8 with a sharp scalpel blade. The T5-T7 spinal column segment was cryoprotected with a sucrose/OCT gradient, 10% sucrose in PBS for 30 minutes, 15% sucrose in PBS for 1 hour, 20% sucrose in PBS for 2 hours, 30% sucrose on PBS for 2 hours, half 30% sucrose/half OCT overnight at 4° C with gentle agitation. Samples were mounted in OCT and rapidly frozen in methylbutane cooled on dry ice. 20 μm sections were cut and mounted on slides, collecting every third section, and allowed to dry for 2 hours at room temperature before freezing or processing for AP staining then immunohistochemistry as described below.

### Section Alkaline Phosphatase Staining

Slide mounted cryosections were washed 3 times for 10 minutes in PBS at room temperature, then were incubated for 30 minutes in 1 mM MgCl2 at 70° C. Sections were moved to a humidified slide staining chamber and were stained with 0.1 M Tris/50 mM MgCl2/0.1 M NaCl in water, pH 9.5 with 16.6 µg/mL BCIP and 33.3 µg/mL NBT and 1.25 mM Levamisole until sufficient staining had developed, 4-48 hours. Once sufficient staining had developed sections were incubated for 15 minutes in 1.25 mM levamisole, 0.25 M EDTA in PBS to stop the AP reaction. Sections were then coverslipped with Fluoromount-G or were processed for fluorescence immunohistochemistry.

### Section Immunohistochemistry

Slide mounted cryosections were washed 3 times for 10 minutes in PBS at room temperature, then were incubated for 30 minutes in blocking solution (5% v/v Normal Donkey Serum, 5% v/v DMSO, 0.25% v/v Triton X-100 in PBS). Slides were incubated in primary antibodies diluted in blocking solution at the concentrations listed in Table 4 overnight at 4° C in a humidified staining box. Slides were washed 3 times for 10 minutes each with PBS, then incubated for 4 hours at room temperature in secondary antibodies diluted at 1:500 in blocking solution. Slides were washed 3 times for 10 minutes each in PBS, adding 1:5000 Hoescht to the first wash. Slides were coverslipped with Fluoromount-G and imaged.

### Skin Section Image quantification

Immunohistochemically labeled and AP-stained skin sections were imaged on a Zeiss AxioImager M.2 microscope with an Apotome.2 structured illumination module. For each animal, a tiled image field approximately 4500 μm wide was captured on a series of 5 nonconsecutive skin sections. The number of hair follicle structures in each section was counted using the DAPI channel. The number of hair follicles with a marker-positive LLE was counted using either the IHC labels described in the text, or AP staining. The proportion of innervated follicles and follicle density measurements from each section were averaged together to generate an average follicle density and follicle innervation proportion value for each animal.

### DRG section Image Quantification

AP stained and immunohistochemically labeled DRG sections were imaged on a Zeiss AxioImager M.2 microscope with an Apotome.2 structured illumination module. Cell counting was performed manually using the Cell Counter plugin in FIJI/ImageJ.

### Statistical Analysis

All statistics were performed in R v4.1.1. Statistical tests, N values, and P-values are described in the text/figure legends where relevant.

## Acknowledgements

We thank the Ginty, Badea, and Nathans Labs for the mouse lines used in the preparation of this manuscript. We would also like to thank the members of the Lumpkin Lab for their help with refining the whole-mount skin immunohistochemistry protocol. Finally, we would like to thank the members of the Wright Lab, Murthy Lab, and Kerstein lab for their feedback on the experiments presented in this manuscript.

## Competing Interests

No competing interests declared

## Funding

This work was supported by the National Institutes of Health [R01NS091027 to K.M.W., T32GM071338 to M.B.P.] and a Whitehall Foundation Research Grant [to K.M.W.].

## Data Availability

Data used in the preparation of this manuscript will be made available upon request.

## Supplemental Figures

**Supplemental Figure 1.**
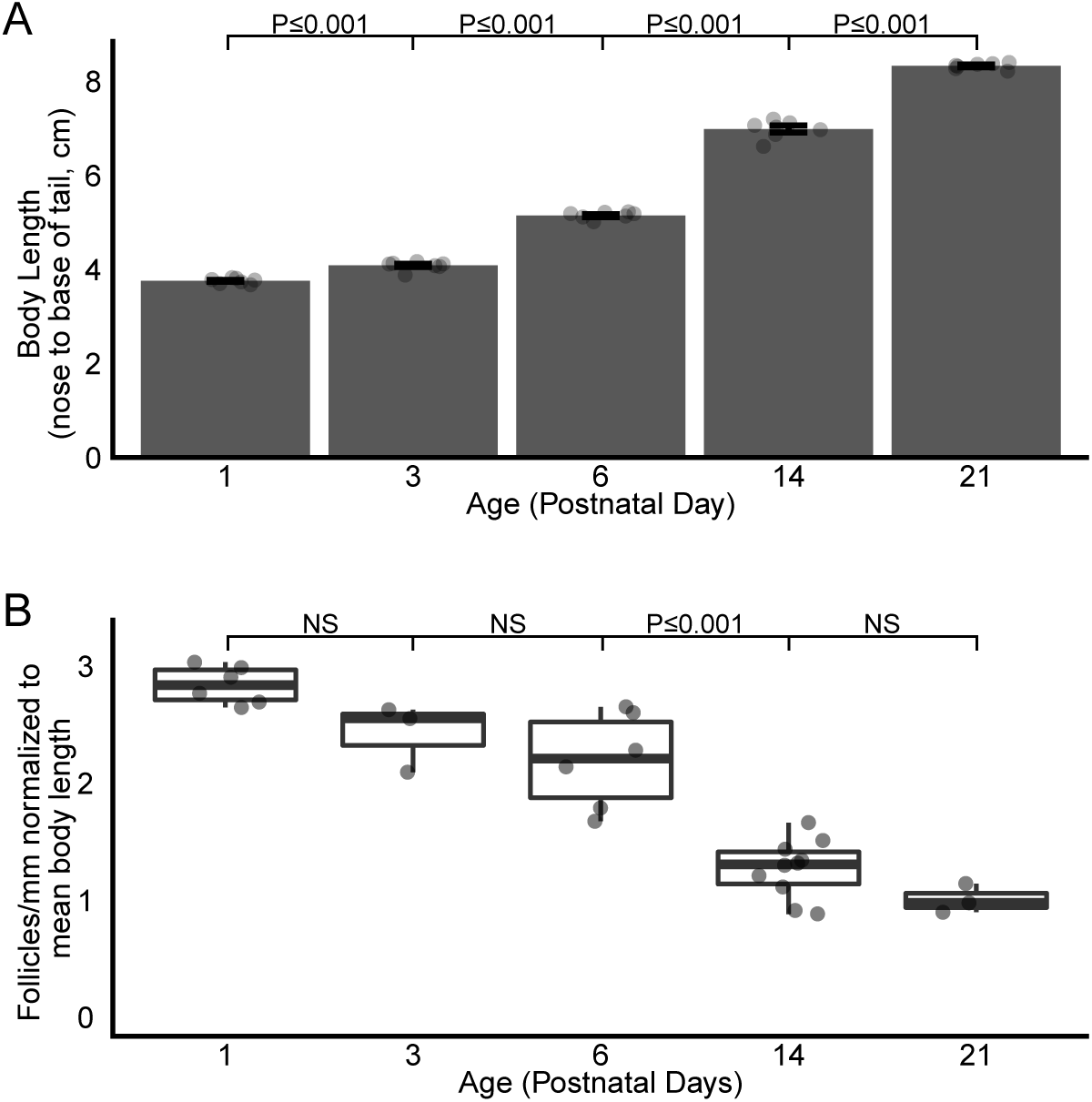
Hair follicle density decreases relative to body length as mice grow. A) Average body length (nose to base of tail) of an age-matched cohort of 10 wild-type mice in centimeters. Bars represent mean ± s.e.m. One-way ANOVA with Tukey’s HSD post-hoc testing (ANOVA P≤ 0.001, P1-P3: P≤ 0.001, P3-P6: P≤ 0.001, P6-P14: P≤ 0.001, P14-P21: P≤0.001. B) Quantification of mouse hair follicle density from Figure 1H normalized to the mean body length of a cohort of 10 same-age mice shown in (A). Bars represent mean ± s.e.m. One-way ANOVA with Tukey’s HSD post-hoc testing (ANOVA P≤ 0.001, P6-P14 P≤ 0.001) N = 6(P1), 3(P3), 6(P6), 10(P14), 3(P21), 5(P60)

**Supplemental Figure 2.**
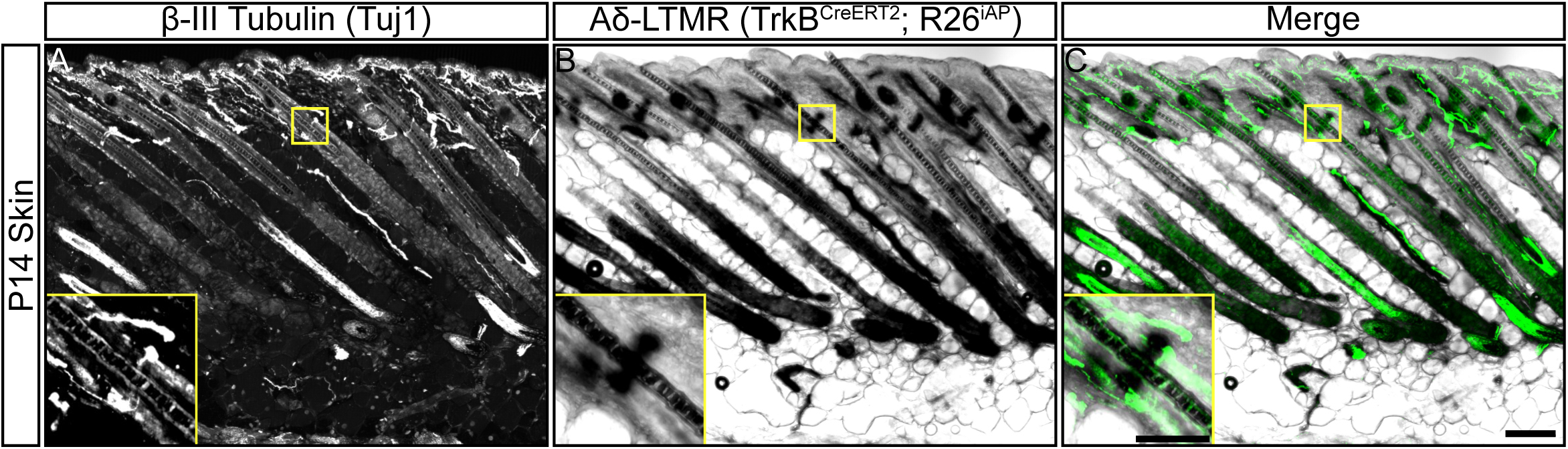
A δ-LTMR neurons fully innervate hair follicles by P14. A-C) Representative image of sagittal plane skin sections from a P14 mouse labeled with markers for all neurons (β-III Tubulin, A), Aδ-LTMRs (*TrkB^CreERT2^;iAP*), B), and merged channels (C). Insets – Higher magnification images of a Tuj1^+^/TrkB^CreERT2+^ double positive longitudinal lanceolate ending. 92.2% ± 3% of follicles were innervated by a TrkB^+^/Tuj1^+^ LLE. This is similar to previously reported values. N = 4 animals. Scale bar – 100 μm, 50 μm inset.

**Supplemental Figure 3.**
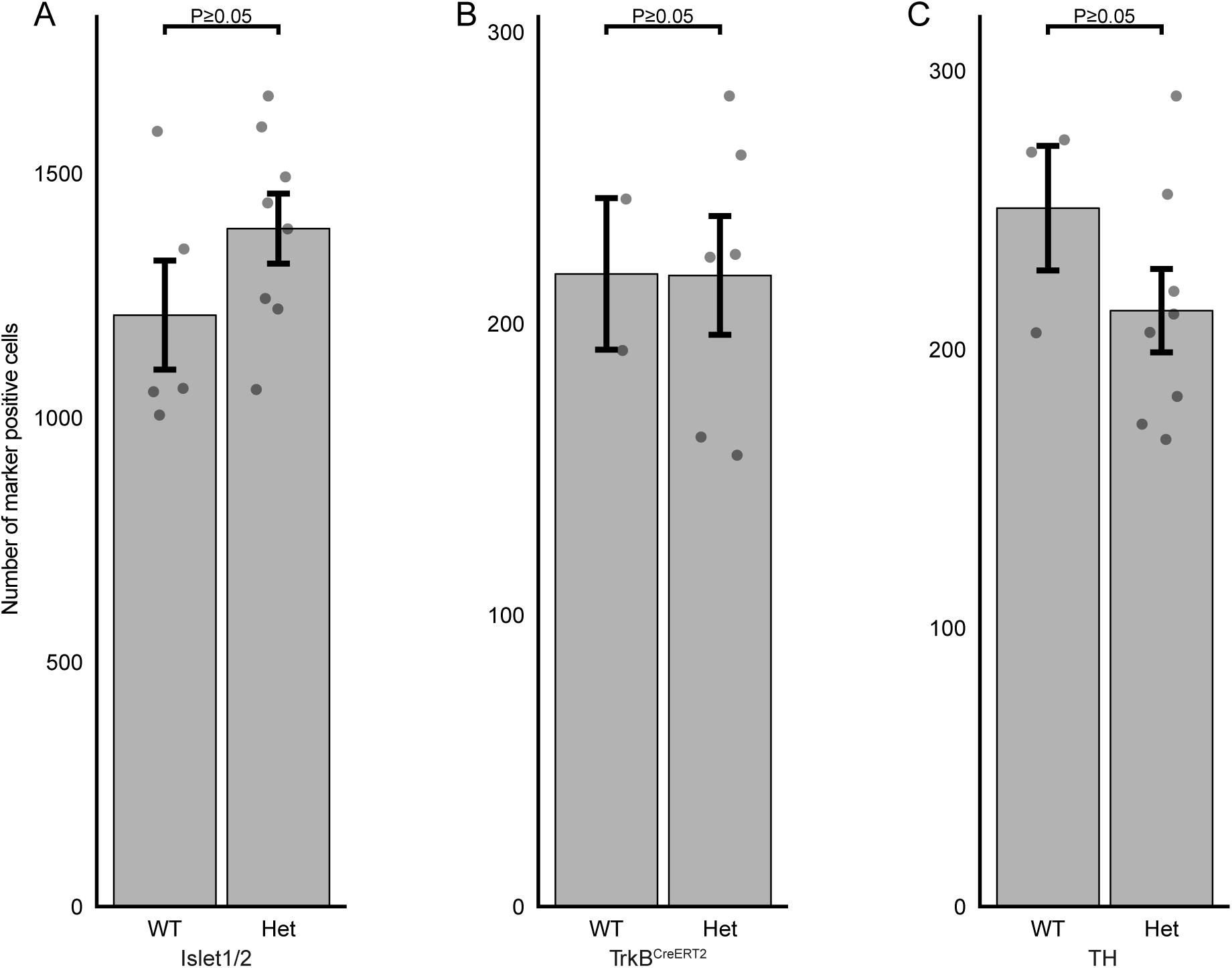
Mice with heterozygous *Bax* deletion have no significant change in the number of neuronal nuclei, Aδ-LTMRs or C-LTMRs compared to wild-type mice. A) Comparison of Islet 1/2^+^ nuclei in *Bax* wild-type and heterozygous mice shows no significant difference. Bars represent mean ± s.e.m. N = 5(WT), 8(Het). Wilcoxon rank sum P≥ 0.05 B) Comparison of TrkB^CreERT2+^ cell bodies in *Bax* wild-type and heterozygous mice shows no significant difference. Bars represent mean ± s.e.m. N = 2(WT), 6(Het). Wilcoxon rank sum P≥ 0.05 C) Comparison of TH^+^ cell bodies in *Bax* wild-type and heterozygous mice shows no significant difference. Bars represent mean ± s.e.m. N = 3(WT), 8(Het). Wilcoxon rank sum P≥ 0.05

**Supplemental Figure 4.**
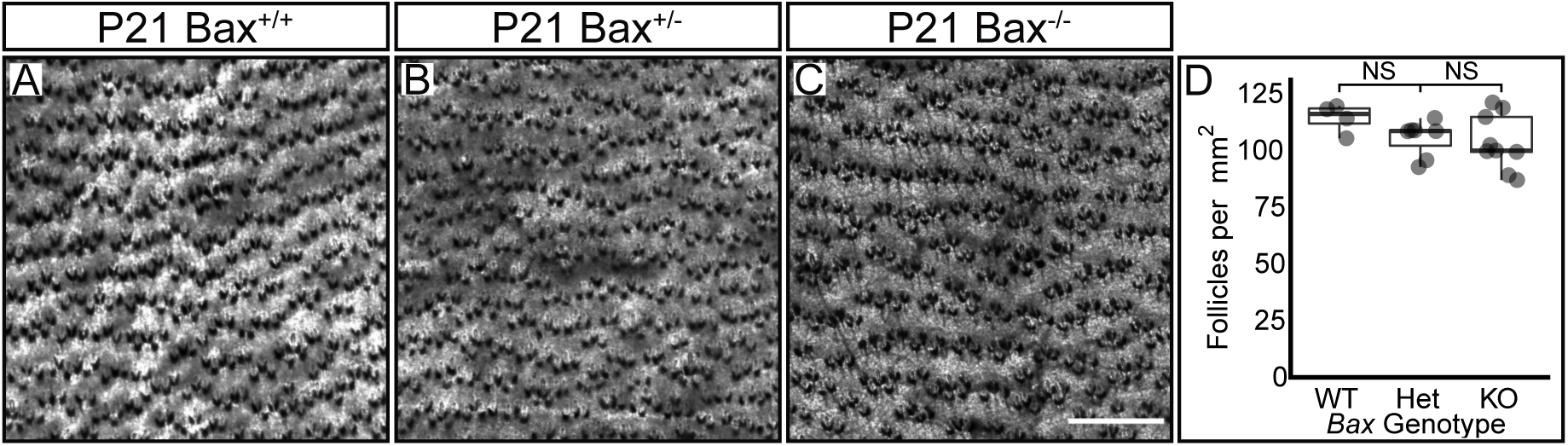
*Bax* deficiency does not change hair follicle density. A-C) Representative images of P21 mouse back skin from *Bax* wild-type (A), heterozygous (B), and knockout (C) animals stained with the lipophilic marker Oil Red O to identify hair follicles. Scale bar – 500 μm D) Quantification of the numbers of hair follicles per square millimeter of back skin. Wild-type: 113.2 ± 3.2 s.e.m. follicles/mm^2^, Heterozygous: 104.3 ± 3 s.e.m. follicles/mm^2^, Knockout: 102.5 ± 4.1 s.e.m. follicles/mm^2^. N = 4(WT), 7(Het), 9(KO). One-way ANOVA P= 0.22

**Supplemental Figure 5.**
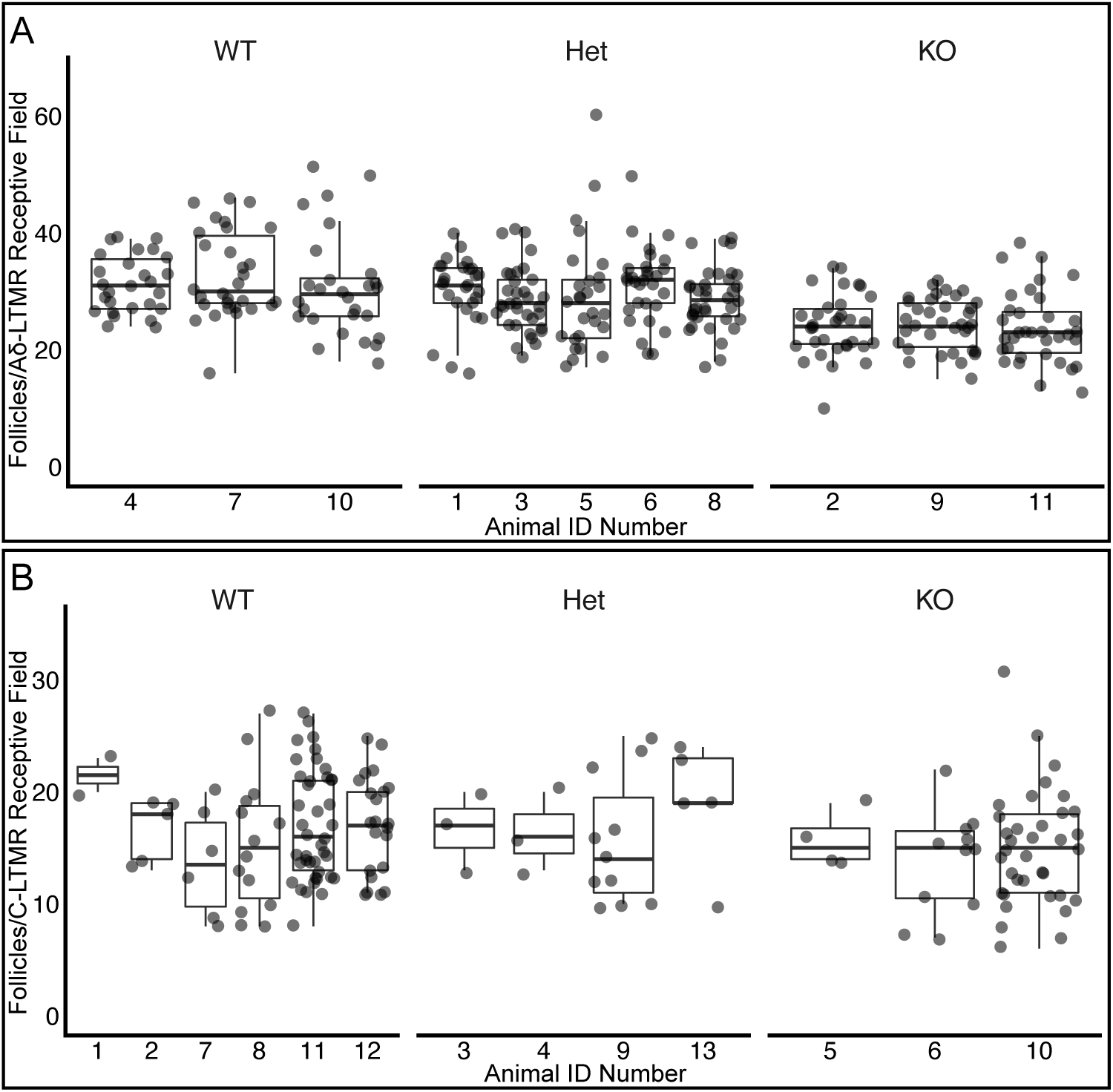
Lack of inter-animal variation in follicles innervated per receptive field between same-genotype animals. A) Quantification of innervated hair follicles per Aδ-LTMR receptive field separated by animal. Each dot represents one quantified receptive field. No significant differences in number of innervated follicles per receptive field were seen between animals within genotype groups. Statistical analysis – One-way ANOVA Bax WT: P= 0.52, Bax Het: P= 0.49, Bax KO: P= 0.86 B) Quantification of innervated hair follicles per C-LTMR receptive field separated by animal. Each dot represents one quantified receptive field. No significant differences in number of innervated follicles per receptive field were seen between animals within genotype groups. Statistical analysis – One-way ANOVA Bax WT: P= 0.38, Bax Het: P= 0.70, Bax KO: P= 0.72

**Supplementary Table 1:** Summary follicle innervation data shown in **Figures 1-3**.

| Age | Follicles/mm<br>± s.e.m.<br>(Fig. 1B-G) | % Tuj1<sup>+</sup><br>± s.e.m.<br>(Fig. 2A-F) | % Calbindin<sup>+</sup><br>± s.e.m.<br>(Fig. 3A-F) | % TH<sup>+</sup><br>± s.e.m.<br>(Fig. 3H-M) | % TrkB<sup>+</sup>/Tuj1<sup>+</sup> ±<br>s.e.m.<br>(Supp. Fig. 2A-C) |
| --- | --- | --- | --- | --- | --- |
| P1 | 10.66 ± 0.25 | 4.35 ± 1.28 | 0.05 ± 0.05 | 3.64 ± 0.59 |  |
| P3 | 9.90 ± 0.68 | 54.44 ± 1.50 | 20.16 ± 1.31 | 52.52 ± 2.03 |  |
| P6 | 11.25 ± 0.85 | 83.68 ± 6.17 | 15.44 ± 1.46 | 92.54 ± 1.26 |  |
| P14 | 8.85 ± 0.55 | 94.99 ± 0.94 | 17.07 ± 2.39 | 94.88 ± 1.07 | 92.2 ± 3.00 |
| P21 | 8.37 ± 0.60 | 94.33 ± 0.95 | 19.30 ± 2.95 | 94.15 ± 1.77 |  |
| P60 | 5.04 ± 0.48 | 90.69 ± 2.88 | 10.59 ± 1.83 | 97.61 ± 1.20 |  |

**Supplementary Table 2:**
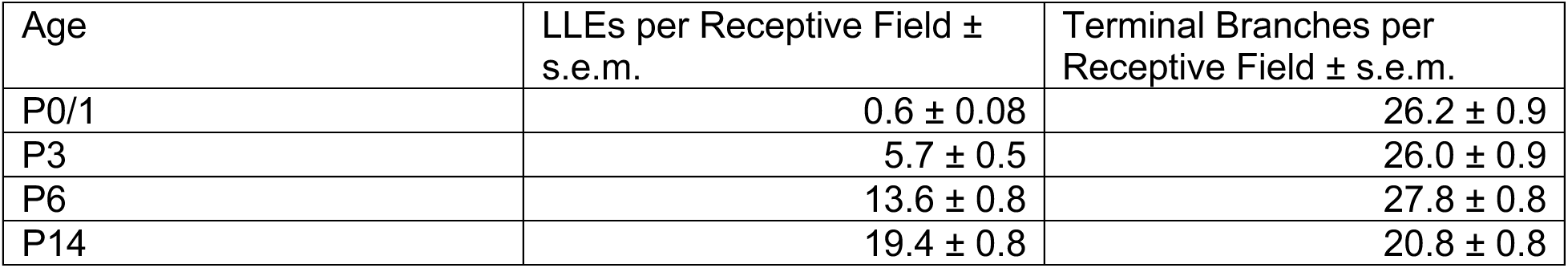
Summary Aδ-LTMR receptive field development data shown in **Figure 4**.

| Age | LLEs per Receptive Field ±<br>s.e.m. | Terminal Branches per<br>Receptive Field ± s.e.m. |
| --- | --- | --- |
| P0/1 | 0.6 ± 0.08 | 26.2 ± 0.9 |
| P3 | 5.7 ± 0.5 | 26.0 ± 0.9 |
| P6 | 13.6 ± 0.8 | 27.8 ± 0.8 |
| P14 | 19.4 ± 0.8 | 20.8 ± 0.8 |

**Supplementary Table 3:** Summary of Bax knockout DRG cell count data shown in **Figure 5**.

| Cell Type | Marker | WT/Het Mean Cell Count ± s.e.m. | KO Mean Cell Count ± s.e.m. | KO normalized to WT/Het mean ± s.e.m. |
| --- | --- | --- | --- | --- |
| Neurons | Islet 1/2 | 1318.8 ± 63.6 | 1573.4 ± 112.6 | 1.19 ± 0.09 |
| Aδ-LTMR | TrkB<sup>CreERT2</sup> | 216.6 ± 15.7 | 332.0 ± 3.0 | 1.53 ± 0.01 |
| C-LTMR | TH | 223.9 ± 12.9 | 300.0 ± 39.7 | 1.34 ± 0.18 |

## Notes

### Competing Interest Statement

The authors have declared no competing interest.

